# Intraepithelial lymphocytes exhibit selective immunity to intestinal pathogens

**DOI:** 10.1101/2025.05.14.654011

**Authors:** Amanpreet Singh Chawla, Olivia J. James, Purbasha Bhattacharya, Dina Dikovskaya, Suzanne L. Hodge, Maud Vandereyken, Lee Robinson, Henry M McSorley, Mattie C Pawlowic, Mahima Swamy

**Author notes:** Address correspondence to: Dr. Mahima Swamy.

## Abstract

Intraepithelial lymphocytes (IEL) are abundant, tissue-resident T cells critical for intestinal immune surveillance, yet their precise roles have remained elusive due to the lack of models enabling their selective genetic ablation. Here, we report the generation of *Gzmb*-Cre knock-in mice, that when crossed with inducible Diphtheria toxin receptor mice, allows targeted and inducible ablation of IELs without perturbing peripheral immunity. Using this model (referred to as IEL^iDTR^), we demonstrate that IELs are dispensable for intestinal homeostasis, including epithelial architecture, barrier function and microbial composition. However, loss of IELs led to increased intestinal infection by *Salmonella* Typhimurium and *Cryptosporidium parvum*, but did not affect responses to *Listeria monocytogenes* infection or DSS colitis. Interestingly, IEL deficiency reduced fecundity of the intestinal nematode, *Heligmosomoides polygyrus*, despite unaltered worm burden, suggesting a permissive role in helminth colonization. These findings position IELs as evolutionary-tuned sentinels against co-evolved pathogens that can cause lethal diarrheal diseases, and establish the IEL^iDTR^ mouse as a vital genetic tool for dissecting IEL function in host-pathogen interactions *in vivo*.

## Introduction

Despite the description of multiple small round lymphocytes within the intestinal epithelium over 150 years ago^1^, little is known about the functions of intestinal intraepithelial lymphocytes (IEL). Most IEL (>90%) are T lymphocytes and they are present within the small intestinal epithelium at a rate of 1 for every 10 epithelial cells, thus representing the largest population of T cells in the body^2,3^. They are found in all jawed vertebrates, including in cartilaginous fish, such as sharks and rays, that lack organized lymphoid structures, and are thought to have arisen to provide protective immunity in the context of increased injury and infection in jawed animals^4^. IELs constantly monitor the epithelium for barrier disruption or cellular stress caused by pathogens and mount an appropriate protective response^5,6^. IELs may kill infected or dysregulated cells^7^, enhance repair of damaged epithelium^8^, and develop memory^9^ to maintain intestinal homeostasis. Yet precise definition of the functions of IEL remain unclear^10^.

Comprehensive analysis of IEL biology has been hindered by the absence of mouse models that enable specific targeting of intestinal IELs. This challenge stems from the developmental heterogeneity of IEL populations, which comprise both thymic-derived natural IELs that undergo selection on self-antigens during early development, and induced IELs that originate from conventional peripheral T cells responding to dietary and microbial antigens. Although typically classified into TCRγδ and TCRαβ IEL—together comprising ∼95% of IELs—this framework masks considerable phenotypic diversity. IEL populations comprise natural T cells derived from thymic precursors that seed the gut early in development (TCRαβ^+^CD8αα^+^ or TCRγδ^+^CD8αα^+^ or TCRγδ^+^CD8^-^ cells), as well as induced IELs (TCRαβ^+^CD8αβ^+^CD8αα^+^ or TCRαβ^+^CD4^+^CD8αα^+^) derived from conventional peripheral T cells. The induced IEL lineages include the tissue-resident memory (T_RM_) cells that are generated through viral infections and provide long-lived antigen-specific protection^9^. Natural αβIELs are thought to have a more immunoregulatory function, as adoptive transfer of these cells can prevent induction of T cell mediated colitis^11^, and natural γδIELs can express epithelial growth factors^8,12^. Thus distinct IEL subsets may serve different functions within the intestinal epithelium. However, there are also multiple indications of functional redundancy in IEL populations. All IEL subsets express cytotoxic molecules including granzyme A and granzyme B^7,13^ and proinflammatory cytokines such as IFNγ^14^, and both induced and natural IEL can produce antimicrobial peptides^15^.

Proteomic and single-cell RNA-seq data indicate a high level of similarity between αβIELs and γδIELs^13,16,17^. Indeed, in both human and mouse scRNA-seq datasets, natural αβIELs and γδIELs are indistinguishable except for their TCR genes. αβIELs and γδIELs also display very similar functional responses to IL-15, and in their ability to induce responses in epithelial cells^16,18^. Yet previous studies have mainly assigned functions to IELs by analyzing TCRδ-deficient (*Tcrd-/-)* mice, which only lack γδIELs, or TCRβ-deficient mice, that lack αβIELs as well as all peripheral αβ T cell populations^5,19,20^. As αβIELs still populate the guts of γδ-deficient mice and vice versa, these studies may have underrepresented the importance of IELs. For example, multiple studies have shown that IEL respond to intestinal infection with the bacterial pathogen *Salmonella*^5,6^, yet *Tcrd^-/-^* mice show only transient loss of protection against infection^15^. Therefore, a key goal to better understand the function of IEL in vivo is to generate a mouse model that permits deletion of all IEL subsets, without affecting other T cell subsets.

Here, we developed a new mouse model that enables selective and inducible depletion of all intestinal IEL subsets while preserving systemic T cell populations. This model exploits our observation that Granzyme B (GzmB) is highly expressed in all IEL subsets and is only expressed in IELs *in vivo* under homeostatic conditions. We generated a mouse model expressing Cre recombinase knocked in (KI) to the endogenous *Gzmb* locus, **GzmB^CreKI^**. By crossing the Gzmb^CreKI^ mice to both the Cre-inducible tdTomato reporter and an inducible diphtheria toxin receptor (iDTR) mouse model^21^, we generated a diphtheria toxin (DT)-inducible IEL depletion model, IEL^iDTR^ mice. Studies in these mice reveal that IELs are dispensable for gut homeostasis but that they exhibit selective control of different intestinal pathogens.

## Results

### GzmB homogeneously marks IELs at steady state

A major obstacle in developing IEL-specific mouse models stems from the profound heterogeneity of IEL populations. No markers have yet been identified that specifically labels all IEL and no other immune compartments, other than anatomical location. However the integrin α_E_(CD103)β_7_, that binds to E-cadherin on intestinal epithelial cells, labels >90% of IEL^22^. Therefore we focussed our analyses on CD103+ immune cells isolated from the small intestinal epithelium. High-dimensional flow cytometry of from the mouse small intestinal epithelium delineates at least eight subsets among IEL (delineated as CD45+CD103+): among TCRγδ IELs, the predominant population is CD8αα⁺, with minor CD8αβ⁺ and CD4⁻CD8⁻ populations; TCRαβ IEL include natural CD8αα⁺, induced CD8αβ⁺, and rare subsets such as CD4⁺CD8α⁺ and CD4⁻CD8⁻ IEL (**Fig. 1A and Suppl. Fig 1A)**. Despite this apparent heterogeneity based on surface markers, analysis of proteomic data from three major IEL subsets (TCRαβ CD8αα, TCRγδ CD8αα, TCRαβ CD8αβ) revealed >95% shared proteins (**Fig. 1B**). We therefore hypothesized that IEL subsets might converge on a shared transcriptional identity that is driven by the intestinal microenvironment. To test this, we performed single-cell chromatin accessibility profiling (scATAC-seq) across mouse small intestinal CD8α+ IELs. Notably, we observed marked epigenetic convergence across all IEL populations (**Fig. 1C and Suppl. Fig 1B)**, suggesting a unifying regulatory architecture. We further examined chromatin accessibility at *Itgae* (encoding CD103), a canonical IEL marker. Over 97% of IELs exhibited open chromatin at the *Itgae* locus **(Suppl. Fig. 1C**). Similarly, ∼70% showed accessible chromatin at *Ifng*, consistent with their known Type 1-skewed effector program **(Suppl. Fig. 1D)**. We also reanalyzed previously published bulk ATAC-seq datasets of TCRγδ and TCRαβ mouse IELs^23^. In the bulk ATAC-seq data, IEL subsets shared 82% of accessible peaks (23,699/28,789), underscoring a high degree of epigenomic overlap, and emphasizing molecular convergence across both chromatin and protein levels **(Suppl. Fig. 1E)**. Together, these findings argue against lineage-centric views of IEL and support a model in which both αβ and γδ IEL subsets form a cohesive, functionally unified guardian of the intestinal epithelium.

**Figure 1.**
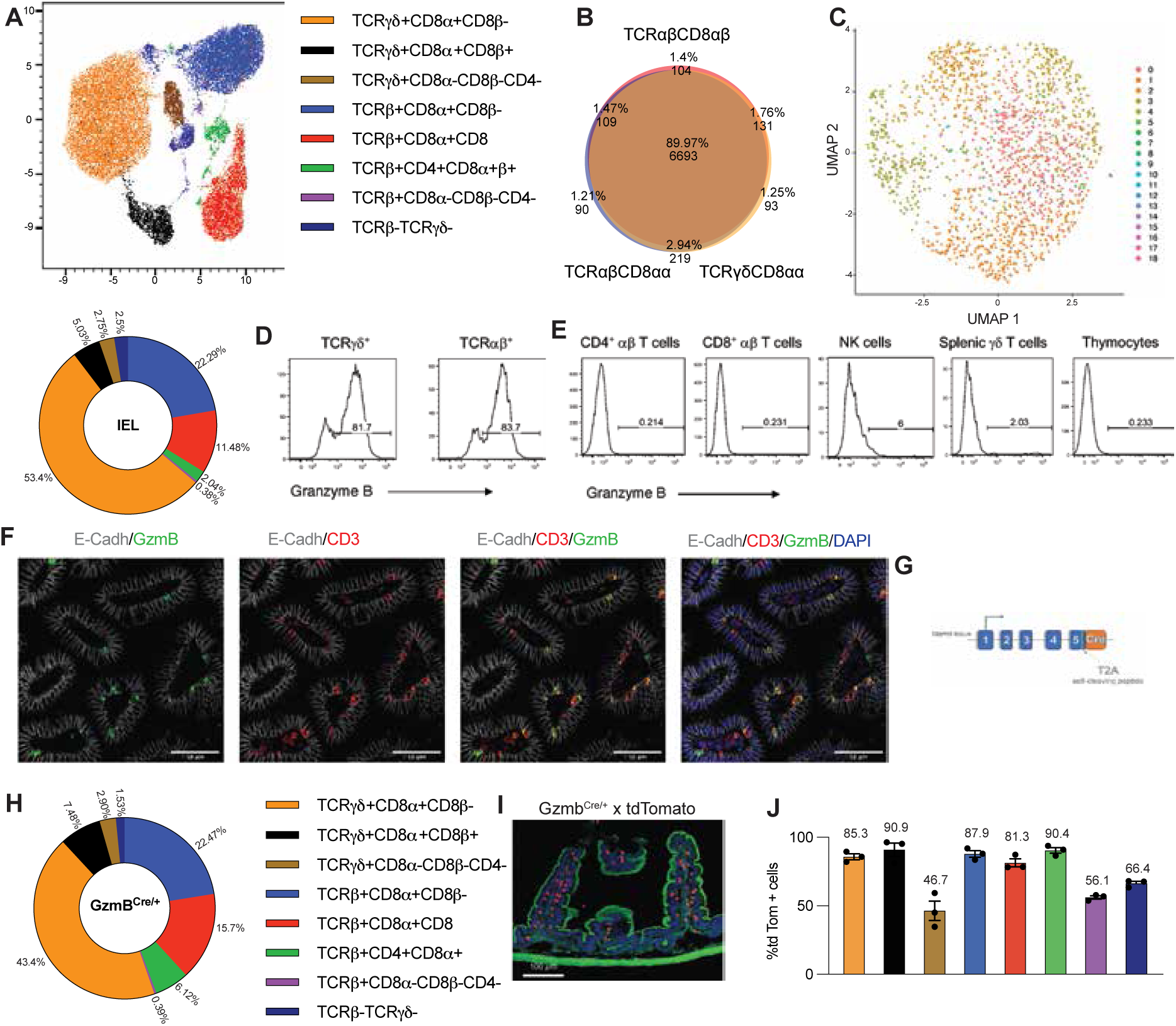
Granzyme B uniformly marks IEL at steady state. **A)** UMAP projection of CD45⁺CD103⁺ IEL isolated from the small intestinal epithelium, revealing phenotypic heterogeneity across subsets. Accompanying pie chart depicts relative frequencies of each subset. Representative of three mice. **B)** Area-proportional Venn diagram illustrating shared proteomic profiles among three major IEL subsets: TCRαβ⁺CD8αα⁺, TCRγδ⁺CD8αα⁺, and TCRαβ⁺CD8αβ⁺. **C)** UMAP embedding of CD45⁺CD3⁺CD8α⁺ IEL profiled by single-cell ATAC-seq, demonstrating convergence in chromatin accessibility landscapes. **D–E)** Representative intracellular flow cytometry histograms of GzmB protein expression in (D) IEL and (E) peripheral immune populations, including NK cells, thymocytes, and splenic T cells. **F)** Confocal images of jejunal tissue sections stained for GzmB (green), CD3 (red), E-cadherin (gray), and nuclei (DAPI, blue), showing GzmB⁺CD3⁺ cells localized to the epithelial compartment. Scale bar, 100 μm. Representative of five mice. **G)** Schematic of the *Gzmb*-Cre knock-in allele, in which a T2A-Cre cassette is inserted downstream of the endogenous *Gzmb* stop codon, enabling bicistronic expression. **H)** Pie chart depicting the relative frequencies of each IEL subset in Gzmb^Cre/+^mice. Representative of three mice. **I)** Representative immunofluorescence image of jejunal sections from Gzmb^Cre/+^× Rosa26^LsL-tdTomato^ mice, showing tdTomato⁺ IELs (red) within the epithelium. Actin (phalloidin, green); nuclei (DAPI, blue). Representative of three mice. **J)** Flow cytometry plots and pooled quantification of tdTomato⁺ frequencies across IEL subsets, with individual bar labels denoting mean percentage of tdTomato⁺ cells within each population. Data represents mean ± SEM from 3 mice/group.

Based on the high level of proteomic and epigenetic similarity between IEL populations, we hypothesized a high amount of functional redundancy. Therefore, we aimed to identify a molecule that would allow us to specifically target all IEL subsets while leaving other immune cells intact. We revisited previously published IEL proteomics datasets and compared them with other immune cell proteomics data from the Immpres database (www.immpres.co.uk^24^), and also analyzed transcriptomic data of mouse immune cells on the Immgen database. Of all the proteins that were enriched in IEL compared to other immune cell subsets, only granzymes (Gzm) emerged as potential candidate molecules. Although there were other molecules found in IEL that were not found in peripheral T cells, most of these molecules were either epithelial molecules or highly expressed by a subset of immune cells, such as natural killer (NK) cells^13^. By contrast, granzyme B (GzmB) is expressed highly in >80% of all IELs (**Fig. 1D**) and is only expressed at low levels in NK cells and not in conventional T cells in the resting state (**Fig. 1E**). Further, unlike GzmA that is expressed during early T cell development (rendering it unsuitable for specific targeting^25^), GzmB is not expressed during thymocyte development (**Fig. 1E**). These data were corroborated by analysis of RNA-seq data on the Immgen database **(Suppl. Fig. 1F)**. *GzmB* mRNA was only detected in NK cells and in IEL in uninfected steady state mice, although its expression was induced in activated NK cells and activated CD8 T cells post infection (marked as post-infection NK cells, **Suppl. Fig. 1F**). However, previous research has shown that in NK cells, only the mRNA of *GzmB* and not the protein is expressed in the resting state^26^, consistent with our flow cytometry data. Moreover, NK/ILC1 cells are very few compared to IEL within the small intestine (**Fig. 1A**, contained within TCRβ-TCRγδ-cells). Indeed, immunostaining of GzmB and CD3 in mouse small intestinal tissue localized GzmB⁺ cells to the epithelial layer, and GzmB+ cells were also CD3+, reinforcing the idea that GzmB is specifically expressed only in IEL in the small intestines of mice (**Fig. 1F**). Taken together, these findings establish GzmB as a definitive marker of mouse IEL at steady state.

Having established *Gzmb* as a potential specific marker for intestinal IELs, we investigated whether Gzmb-based Cre strains could be used to target IEL. However, when we crossed a previously established GzmB-Cre transgenic mouse model^27^, which utilized a truncated human *GZMB* promoter, to a *Rosa26^LSL-EYFP^* reporter, we did not detect labelling of IELs **(Suppl. Fig. 1G)**. Interestingly, previous attempts at targeting NK cells with the same GzmB-Cre transgenic mouse model^28^ showed that only activated NK cells were efficiently targeted. This transgenic Gzmb-Cre was also spuriously active in CD4 T cells and even in hematopoietic stem cells that did not express mouse GzmB, potentially due to the absence of mouse essential cis-regulatory elements^29,30^. To overcome these limitations, we generated a *Gzmb*-Cre knock-in (KI) mouse by inserting Cre recombinase into the endogenous *Gzmb* locus, replacing the stop codon with a self-cleaving T2A peptide followed by Cre to preserve native GzmB expression (**Fig. 1G**). Gzmb^Cre/+^ mice developed normally and had normal numbers and distribution of intestinal IEL (**Fig. 1H**). However, despite the 2A peptide, we noticed a slight decrease in GzmB expression in IELs **(Suppl. Fig. 1H)**. Therefore, to control for any effects that could be attributed to reduced GzmB expression, in all further experiments Gzmb^Cre/+^ mice were used as controls. We next crossed the Gzmb^Cre/+^ mice to Rosa26^lsl-tdTomato^ (ref ^31^) reporter mice, where we observed robust and specific tdTomato expression in *Gzmb*-expressing intestinal IEL (**Fig. 1I**). Flow cytometry confirmed that Gzmb^Cre/+^ efficiently activated tdTomato reporter expression in all IEL subsets (**Fig. 1J**). Importantly, the major IEL subsets (TCRγδ CD8αα (orange), TCRγδ CD8αβ (black), TCRαβ CD8αα (blue), TCRαβ CD8αβ (red)), that together account for >90% of all IELs, are all >90% tdTomato positive. Thus, the newly established Gzmb^Cre/+^ effectively labels and targets all IEL subsets for genetic modulation.

### A Gzmb^Cre^ deleter strain enables specific and inducible depletion of IELs

To develop a model that would allow us to inducibly ablate IELs, we crossed Gzmb^Cre/+^ mice to the inducible Diphtheria Toxin Receptor (iDTR), Rosa26^LSL-iDTR^ (ref^21^) mice, enabling diphtheria toxin (DT)-induced ablation of GzmB-expressing cells. From this point onwards we refer to Gzmb^Cre/+^ and Gzmb^Cre/+^;Rosa26^LSL-iDTR^ strains as IEL^WT^ and IEL^iDTR^, respectively. Just a single intraperitoneal injection of DT led to a marked depletion of CD3⁺ IELs identified within the epithelium in small intestinal tissue sections (**Fig. 2A**). However, in the lamina propria, CD3+ T cells were still visible. Flow cytometry analysis confirmed an 80-90% reduction of only CD45+ CD103+ cells in the small intestinal epithelial layer, i.e., IEL, but not CD45+ CD103-cells which includes non-tissue-resident cells (**Fig. 2B**). DT treatment had no effect on Gzmb^Cre/+^ mice without iDTR expression **(Suppl. Fig 2A)**. Further analysis confirmed that DT treatment ablated all IEL subsets by at least 80% (**Fig. 2C-D, Suppl. Fig. 2B)** with γδIELs reduced by 83% and αβIELs reduced by ∼90%. Importantly, DT treatment largely spared immune populations in the small intestinal lamina propria, colon, secondary lymphoid tissues, and thymus, confirming both the specificity and spatial restriction of the system (**Fig. 2C, 2E and Suppl. Fig 2C-2G**). The absence of thymic alterations further indicates that *Gzmb*-driven Cre activity targets post-thymic IEL populations. Thus, this system enables selective and efficient ablation of IELs in vivo.

**Figure 2.**
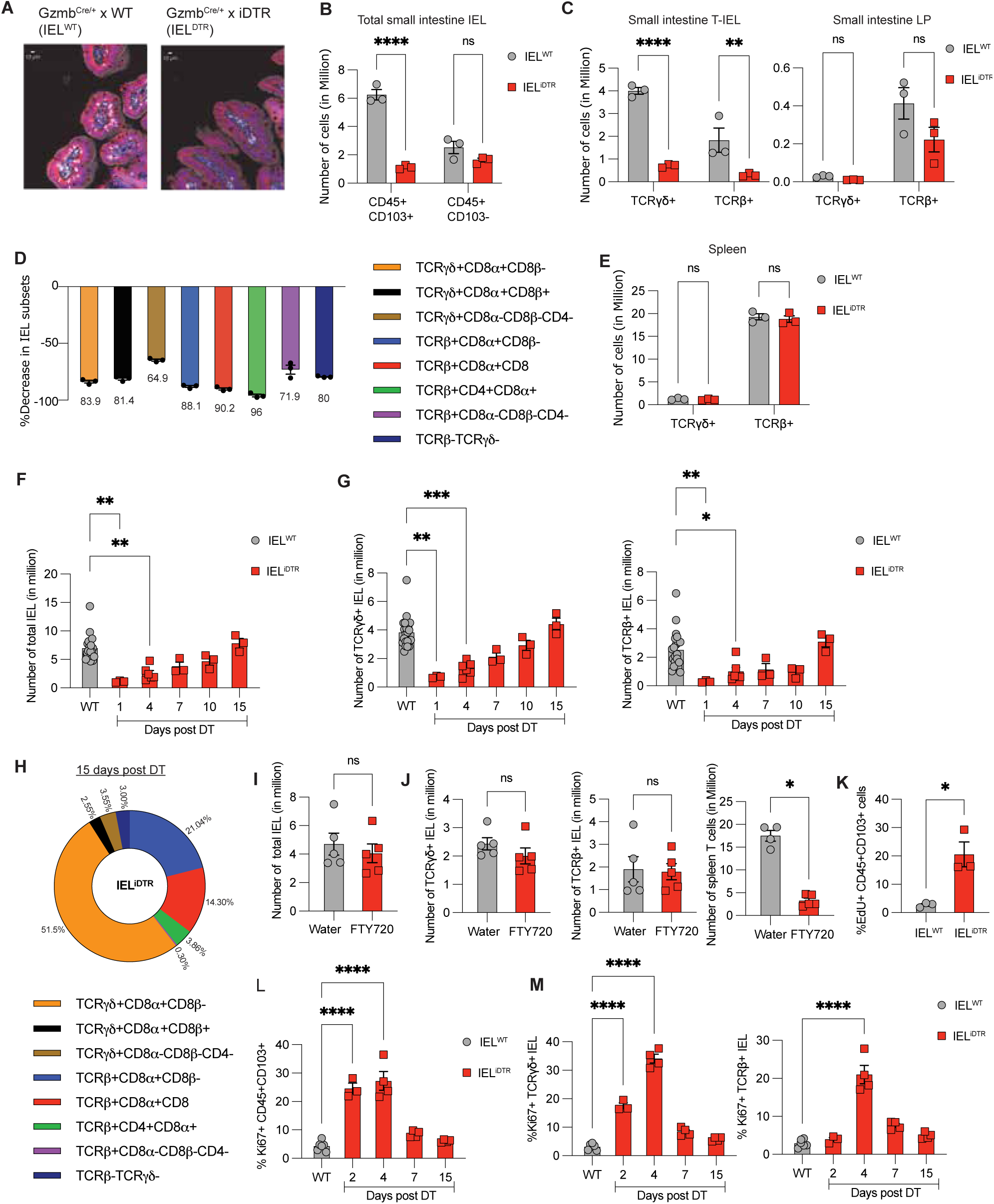
A GzmB-Cre knock-in enables acute and specific IEL ablation. **A)** Representative immunofluorescence image of jejunal tissue from IEL^iDTR^ (*G*zmb^Cre/+^× Rosa26^LsL-iDTR^) and control IEL^WT^ (Gzmb^Cre/+^) mice 24 hours following a single intraperitoneal injection of diphtheria toxin (DT, 50 μg/kg), showing loss of CD3⁺ IELs in the epithelium. Scale bar, 100 μm. Representative of 5 mice. **B)** Flow cytometric quantification of CD45⁺CD103⁺ IELs and CD45⁺CD103⁻ cells in the epithelial fraction 24 h post-DT. Data represents mean ± SEM from 3 mice/group. **C)** Quantification of TCRγδ⁺ and TCRβ⁺ cells by flow cytometry in both IEL and lamina propria (gated on CD45+CD103-) (LP) compartments of the small intestine 24 h post-DT. Data represents mean ± SEM from 3 mice/group. **D)** Percent decrease across individual IEL subsets in IEL^iDTR^ mice 24 h after DT, compared to IEL^WT^ with individual bar labels denoting mean percentage of cells deleted within each population. Data represents mean ± SEM from 3 mice/group. **E)** Flow cytometric analysis of TCRγδ⁺ and TCRβ⁺ T cell populations in the spleen 24 h post-DT administration. Data represents mean ± SEM from 3 mice/group. **F–G)** Time-course quantification of total IELs (CD45⁺CD103⁺), TCRγδ⁺, and TCRαβ⁺ subsets at multiple time points post-DT (days 1, 4, 7, 10, 15), showing near-complete repopulation by day 15. Data represents mean ± SEM from 3 mice/group. **H)** Distribution of IEL subsets 15 days following DT treatment, demonstrating restoration of pre-depletion composition. Representation of 3 mice/group. **I–J)** Quantification of IEL and spleen T cells recovery at day 7 post-DT in mice treated with FTY720 (to block thymic egress) versus vehicle, demonstrating thymus-independent regeneration. Data represents mean ± SEM from 5 mice/group. **K)** EdU incorporation in CD45⁺CD103⁺ IEL on day 4 post-DT, indicating robust local proliferation during repopulation. Data represents mean ± SEM from 5 mice/group. **L–M)** Ki67 expression in IEL across time points post-depletion, showing a transient spike in proliferation that returns to baseline by day 15. Data represents mean ± SEM from 4-6 mice/group. Statistical analyses: ANOVA with Sidak’s post hoc correction (B,C,E, F, G, L, M); unpaired t-test (I, J, K). **\*** p < 0.05, ** p < 0.01, *** p < 0.001, **** p < 0.0001, ns = non-significant.

### Rapid in situ proliferation drives IEL compartment reconstitution

Having established that our Gzmb^Cre/+^ model enables efficient and specific intestinal IEL ablation, we next sought to determine whether the IEL compartment can regenerate post-deletion and, if so, how rapidly. We depleted IELs and harvested mice at day 1, 4, 7, 10, and 15 post-DT treatment. Surprisingly, the CD45+ CD103+ cells in the IEL compartment were ∼50% replenished by day 7 and fully restored by day 15 (**Fig. 2F**), with total IEL numbers and subset frequencies indistinguishable from wild-type controls (**Fig. 2G-H and Suppl Fig. 3A-B)**. This rapid recovery was unexpected, as the conventional view is that tissue-resident T cells are stable, long-lived populations that therefore do not require active replenishment ^32^.

This raised the broader question of whether such rapid turnover is an intrinsic feature of IEL biology at steady state. To test this, we used the previously reported Gzmb^CreERT2^ model^29^ crossed to *Rosa26^lsl-tdTomato^* reporter mice, allowing us to time-stamp label IELs at a defined zero-time point following tamoxifen treatment and monitor their subsequent loss over time. The labelling efficiency of IELs in Gzmb^CreERT2^ x Rosa26^LsL-tdTomato^ reporter mice treated with 4-hydroxytamoxifen was around 65% **(Suppl Fig. 3C-3D**). These data show that the transgenic Gzmb^CreERT2^ mouse line, which used a bacterial artificial chromosome (BAC) transgene that included the whole *Gzmb* locus to control Cre-ERT2 expression, worked effectively to inducibly label IEL. We analyzed the Gzmb^CreERT2^ x Rosa26^LsL-tdTomato^ reporter mice 7, 30 or 60 days after 4-hydroxytamoxifen induction of tdTomato expression. Indeed, tdTomato+ IEL exhibited a half-life of >60 days, consistent across TCRγδ+ and TCRαβ+ subsets **(Suppl Fig. 3E),** surpassing previous estimates of 10-36 days based on bromodeoxyuridine labeling experiments^32^. This suggested that IELs are indeed long-lived cells, contrasting with the rapid restoration of the IEL compartment in our Gzmb^Cre/+^ model. Thus, the rapid reconstitution of the IEL compartment in IEL^iDTR^ mice is most likely due to lymphopenia in the niche.

**Figure 3.**
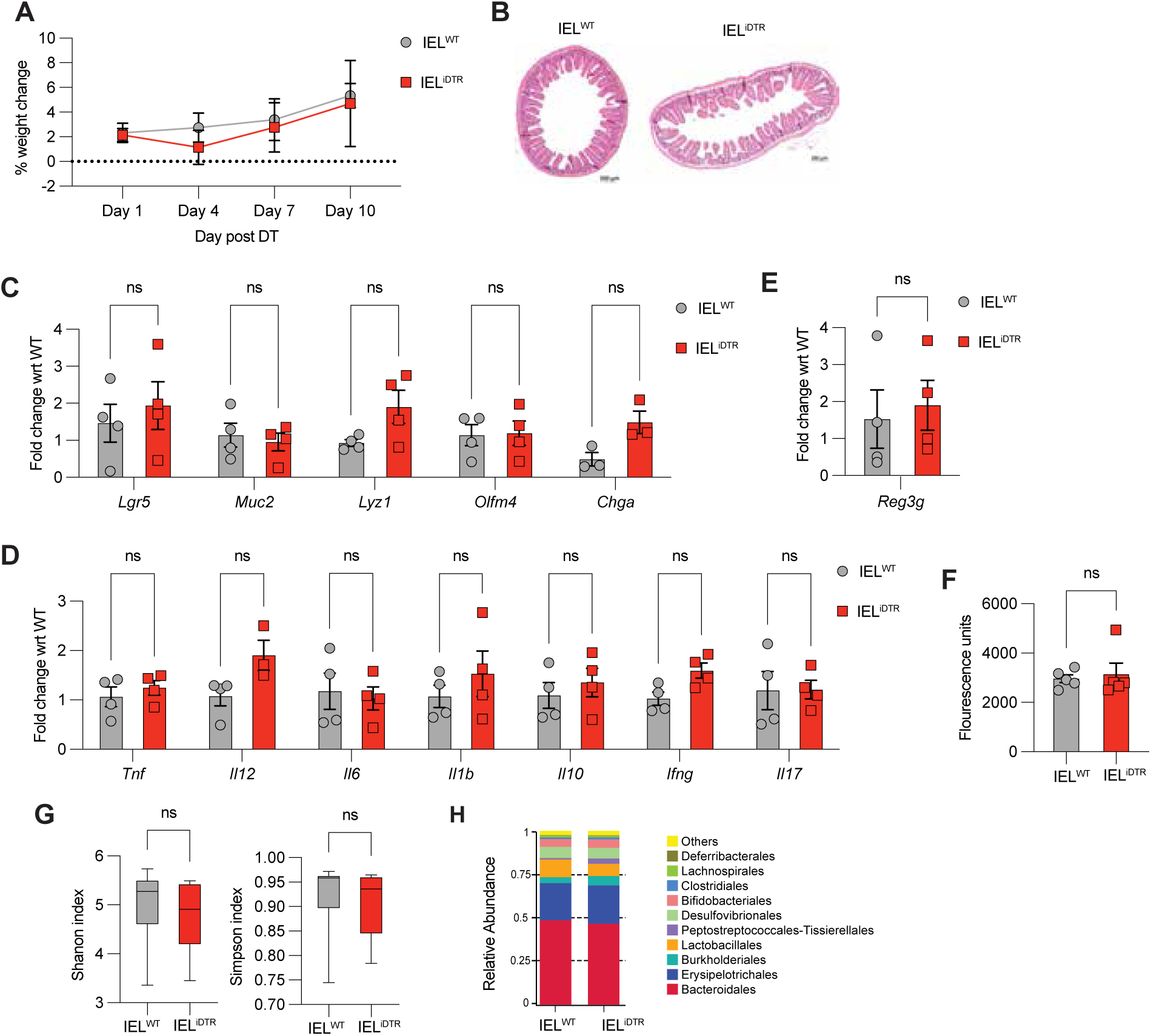
IEL loss does not disrupt intestinal homeostasis at steady state. **A)** Body weight trajectory in IEL^WT^ and IEL^iDTR^ mice over 48 h following DT administration, showing no differences post-IEL ablation. Data represents mean ± SEM from 3 mice/group. **B)** Representative H&E-stained sections of jejunum from IEL^WT^ and IEL^iDTR^ mice 24 h post-DT, demonstrating preserved epithelial architecture. Scale bar, 200 μm. Representative of 5 mice. **C–E)** RT-qPCR analysis of jejunal tissue at 48 h post-DT showing no significant changes in (C) epithelial differentiation markers, (D) pro-inflammatory cytokines, or (E) the antimicrobial peptide Reg3γ. Data represents mean ± SEM from 4 mice/group. **F)** FITC-dextran permeability assay at 24 h post-DT showing intact intestinal barrier function in IEL^WT^ and IEL^iDTR^ mice. Data represents mean ± SEM from 4 mice/group. **G–H)** 16S rRNA sequencing of small intestinal luminal contents at 48 h post-DT. (G) Relative microbial composition and (H) alpha diversity (Shannon and Simpson indices) reveal no major disruption in microbiota structure following IEL depletion. Data pooled from 10 mice per group. Statistical analyses: ANOVA with Sidak’s post hoc correction (C–F); unpaired t-test (G–H). ns = non-significant.

We considered two main mechanisms of IEL regeneration: (1) thymic-derived progenitors replenish the IEL pool, or (2) in situ proliferation of residual IELs drives compartment reconstitution. To distinguish between these scenarios, we treated mice with FTY720—an inhibitor of lymphocyte egress from the thymus—immediately following IEL ablation. IEL^iDTR^ mice received FTY720 or vehicle for six consecutive days post-DT, and IELs were analyzed on day 7. If progenitors from the thymus were driving repopulation, FTY720 treatment should reduce IEL recovery. However, we found no significant difference in IEL reconstitution between vehicle- and FTY720-treated mice (**Fig. 2I**), even though the splenic compartment had reduced T cells (**Fig. 2J**), confirming that FTY720 had worked. Thus, thymic progenitors are not the primary source of replenishment.

To directly assess whether *in situ* proliferation contributed to IEL compartment restoration, we performed an assay for incorporation of EdU—a thymidine analog that labels actively proliferating cells. IELs isolated on day 4 post-deletion showed significantly increased EdU incorporation compared to controls, indicating active *in situ* proliferation as a mechanism of reconstitution (**Fig. 2K**). To corroborate this, we assessed Ki67 expression by flow cytometry at multiple time points post-DT administration. Ki67⁺ IEL frequencies peaked at day 4 and returned to baseline by day 15 (**Fig. 2L–M**). Together, these findings indicate that the IEL compartment rapidly regenerates via *in situ* proliferation, rather than through recruitment of thymic progenitors. This temporal insight is critical for the design of functional studies, as it defines the window of IEL depletion and regeneration. Overall, our data indicate that IEL are normally stable, long-lived cells, but contain within them progenitors capable of self-renewal to rapidly repopulate the epithelium in case of lEL loss. This ability to rapidly proliferate may also be important to allow rapid IEL responses to intestinal stress or inflammation.

### IEL depletion does not disrupt intestinal homeostasis

To determine whether IEL contribute to basal intestinal homeostasis, we assessed a range of physiological and immune parameters following their depletion. IEL^iDTR^ mice exhibited no changes in body weight or overt histological alterations in small intestinal architecture (**Fig. 3A–B**). Gene expression analysis of the jejunal region of the small intestine showed no significant changes in markers of epithelial differentiation (**Fig. 3C**) or in *Reg3γ*, an antimicrobial peptide that is produced by IEL and epithelial cells^15^ (**Fig. 3D**). We next evaluated whether the acute loss of IEL triggered inflammation, either due to apoptotic cell clearance or the absence of immunoregulatory cues. However, inflammatory cytokine expression remained unaltered (**Fig. 3E**). Epithelial barrier integrity, measured by FITC-dextran permeability, was also unaffected (**Fig. 3F**). Finally, 16S rRNA sequencing of small intestinal luminal contents revealed no significant shifts in microbiota composition or diversity (**Fig. 3G–H**). Collectively, these results indicate that IELs are not essential for maintaining steady-state epithelial architecture, barrier function, or microbial homeostasis.

### IELs confer context-specific protection during intestinal infection and inflammation

Given their dispensability at steady state, we next explored IEL function under diverse intestinal stress conditions. We first assessed the role of IELs in colonic inflammation using the dextran sodium sulfate (DSS) model of colitis. Although previous studies suggested a protective role for IEL in this context^17,33^, they often relied on whole-body knockouts or pan-T cell depletion models, making it difficult to isolate IEL-specific effects. Moreover, DSS-induced injury primarily affects the colon, yet the majority of IEL are found in the small intestine. In DT-treated IEL^iDTR^ mice, where only small intestinal IELs are ablated, IEL depletion had no effect on body weight, disease scores, or histological damage following DSS exposure (**Fig. 4A–C**). These data argue against distinct roles for small intestinal IELs in protecting against acute DSS-induced colonic inflammation.

**Figure 4.**
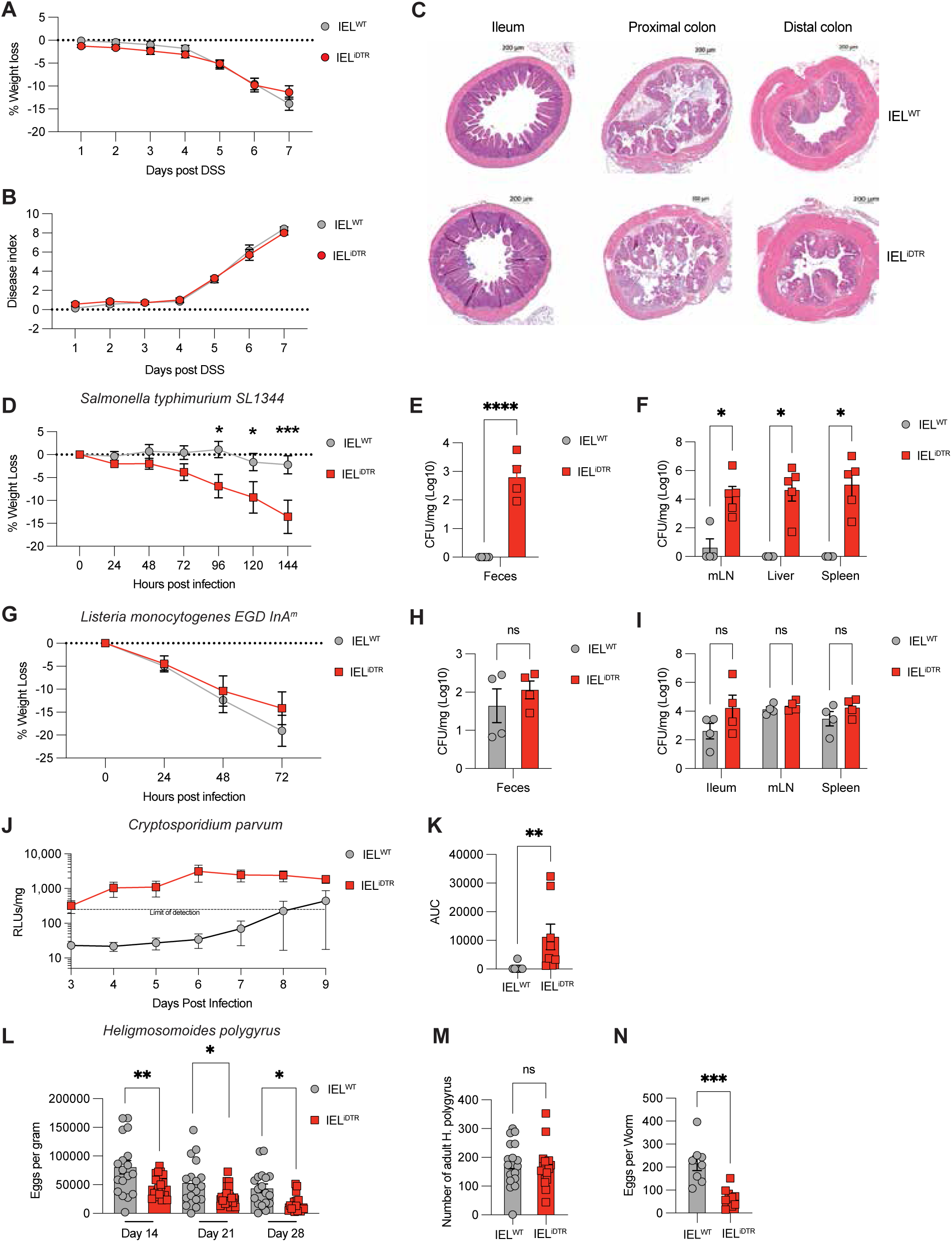
IEL exhibit context-specific roles across diverse infection and injury models. **A–C**) DSS-induced colitis: (A) Body weight change, (B) clinical disease scores, and (C) colon length on day 6 post 3% DSS administration in IEL^WT^ and IEL^iDTR^ mice. Data represent mean ± SEM from 6 mice per group. **D–F)** *Salmonella typhimurium* infection: (D) Body weight loss, (E) fecal bacterial burden at day 3, and (F) systemic dissemination (mLN, liver, spleen) at day 5 post-infection in IEL^WT^ and IEL^iDTR^ mice. Data represent mean ± SEM from 4 mice per group; representative of two independent experiments. **G–I)** *Listeria monocytogenes* (EGD-InlAm) infection: (G) Weight loss, (H) fecal bacterial burden, and (I) systemic bacterial dissemination in IEL^WT^ and IEL^iDTR^ mice. Data represent mean ± SEM from 4 mice per group; representative of two independent experiments. **J–K)** *Cryptosporidium parvum* infection: (J) Luminescence-based quantification of parasite burden and (K) area under the curve (AUC) of cumulative parasite load in in IEL^WT^ and IEL^iDTR^ mice. Data represent mean ± SEM from 8 mice per group. **L–M)** *Heligmosomoides polygyrus* infection: (L) Parasite fecundity (eggs per worm) and (M) total egg burden over time in IEL^WT^ and IEL^iDTR^ mice. Data represent mean ± SEM from 18 female mice per group. Statistical analyses: ANOVA with Sidak’s post hoc correction (A, C, D, F, G, I, M); unpaired t-test (B, E, H, J, K, L), * p < 0.05, ** p < 0.01, *** p < 0.001.

We then explored IEL contributions to pathogen defense. Mice were infected one day post-DT injection to capture the immediate effects of IEL loss on early infection stages. Upon oral challenge with *Salmonella* typhimurium, IEL^iDTR^ mice showed increased weight loss and significantly elevated bacterial loads in feces and systemic tissues 7 days after DT injection compared with IEL^WT^ mice (**Fig. 4D–F).** These data reveal a critical role for IEL in early mucosal containment of *S. typhimurium*. By contrast, IEL depletion had no impact on disease course or bacterial burden following oral infection with *Listeria monocytogenes* EGD Lmo-InlA^m^—a murinized strain engineered to infect murine enterocytes (**Fig. 4G–I**)—highlighting that IEL-mediated protection is not universal across enteric pathogens.

To assess IEL roles in responses to protozoan infection, we employed a bioluminescent strain of *Cryptosporidium parvum*, a pathogen that typically fails to infect immunocompetent mice^34^. Strikingly, DT-treated IEL^iDTR^ mice became permissive to infection, with significantly higher and prolonged parasite burdens than IEL^WT^ mice (**Fig. 4J–K**), uncovering an unexpected requirement for IELs in controlling protozoa that infect intestinal epithelial cells. Finally, we tested the importance of the IEL response to the intestinal helminth *Heligmosomoides polygyrus bakeri*. IEL ablation did not alter adult worm burden but led to significantly reduced egg shedding 14, 21, and even 28 days post IEL depletion and infection (**Fig. 4L–M**), even though IEL numbers should be compltey restored by this timepoint. Thus, our data suggest that IELs support helminth fecundity— possibly by shaping a tissue environment that is conducive to parasite reproduction.

Together, these findings reveal that IELs exert context-specific responses to a spectrum of intestinal challenges. While dispensable for steady-state homeostasis, colitis, and *Listeria* infection, IELs mediate early defense against infections by invasive bacteria (*S. typhimurium*) and protozoa and may even facilitate helminth reproductive success, revealing their tissue-specifc roles in intestinal immunity.

## Discussion

Despite their abundance and evolutionary conservation across vertebrates, the functional contribution of IELs to mucosal immunity has remained ambiguous, largely due to the absence of tools that enable precise genetic manipulation^10^. Previous models targeting IELs relied on either broad promoter activity such as CD4-cre or E8i-Cre, or systemic deletions, each confounded by peripheral effects. Here, we address this gap by developing a *GzmB^Cre^* mouse that permits selective targeting of IEL, including natural and induced αβ and γδ IELs, sparing thymic development and other immune compartments. This *GzmB^Cre^* model not only resolves long-standing concerns about peripheral deletion but provides a much-needed genetic toolkit to dissect IEL biology with spatial precision.

Using this model, we found that IEL are dispensable for maintaining epithelial integrity and microbial homeostasis at steady state but exhibit context-specific functions during intestinal infection. The loss of IEL sensitizes mice to oral *S. typhimurium* and *C. parvum* infections – yet has no impact on *Listeria monocytogenes*. In contrast, IEL appear the support the fecundity of the intestinal helminthic parasite, *H. polygyrus,* a parasite found in all wild mouse populations, where it establishes a persistent infection^35^. This dichotomy suggests that IEL may have evolved under selective pressure to counter persistent, co-evolved enteric threats, rather than opportunistic or environmentally emergent pathogens.

From an evolutionary medicine perspective, our findings suggest that IEL have been selectively shaped to counter ancient, co-evolved intestinal pathogens rather than emergent or opportunistic microbes. IEL depletion sensitized mice to Salmonella spp. and Cryptosporidia, both of which are host-adapted pathogens with deep evolutionary roots in human and animal populations. Salmonella is a facultative intracellular bacterium with a long-standing enteric transmission route in mammals and birds, having co-diverged with vertebrate hosts over millions of years36. Likewise, Cryptosporidium species are obligate enteric parasites that have co-evolved with their hosts and rarely cause fatal disease in immunocompetent individuals, indicating an evolutionary balance favoring transmission over lethality37. In contrast, IEL were dispensable in oral infection with Listeria monocytogenes, a foodborne pathogen and environmental saprophyte whose zoonotic potential emerged relatively recently in human history, likely following agricultural practices and domestication38,39. This dichotomy suggests IEL may have evolved under selection pressure to recognize and contain pathogens with persistent mucosal niches, rather than sporadic, environmentally acquired invaders.

Intriguingly, IEL appear to support rather than restrict the fecundity of H. polygyrus, a helminth with a long co-evolutionary history in mammals40. While total worm burden remained unaffected, IEL-deficient mice exhibited a marked reduction in egg output, implying a permissive role for IEL in maintaining a tissue environment conducive to helminth reproduction. This supports the “old friends” hypothesis, which posits that helminths and microbes that we have co-evolved with over millions of years play regulatory roles in immune homeostasis rather than acting solely as pathogens41. Collectively, these data reposition IELs not as general-purpose defenders but as evolutionarily tuned mediators of intestinal resilience, providing pathogen- and context-specific immunity shaped by ancestral exposure patterns.

Importantly the use of the *Gzmb^Cre-KI^* and the *Gzmb^CreERT2^*mouse models, which we show can both be used to target all intestinal IEL, including αβ and γδ IELs, can be used to establish how IEL might recognize pathogenic infections, and also permit the investigation of other functions of IELs. Understanding how IEL respond to important human enteric pathogens, which cause diarrhoeal diseases (salmonellosis, cryptosporidiosis, listeriosis) that lead to substantial loss of life in young children and older and immunocompromised adults, could be beneficial to mucosal vaccine development and for drugs that can improve infection outcomes by boosting IEL responses.

Together, these findings reposition IEL not as generic sentinels of intestinal epithelium, but as evolutionarily attuned, mediators of mucosal immunity to selected enteric pathogens. The Gzmb-Cre knock-in model thus provides an important platform to explore IEL biology in both mechanistic and disease-relevant settings and offers a tool to systematically dissect how IEL contribute to host defense, tolerance, and tissue adaptation.

## Acknowledgements

The project was funded by Wellcome Trust (Sir Henry Dale Fellowship, 206246/Z/17/Z) and the UK Medical Research Council funding to MS (MC_UU_00038/7). OJJ was supported by a Wellcome Trust PhD studentship (215309/Z/19/Z). We would like to thank A. Rennie and R. Clarke for cell sorting and technical assistance, and the Biological resource unit at the University of Dundee for technical help. We are grateful to ActivMotif for discussions about scATAC-seq data. We also acknowledge help of Dr. Gemma Alderton, Biosciedit, for critical reading and editing of the manuscript.

## Author contributions

MS conceptualized and designed the study, ASC and MS curated and analysed data, ASC, OJJ, MV, DD, PB, SLH, LR, MCP, HJM performed experiments and analysed data. ASC and MS prepared figures and wrote the manuscript with input from all co-authors.

## Materials and methods

### Ethics

Mice were bred and maintained with approval by the University of Dundee ethical review committee under a UK Home Office project license (PD4D8EFEF, PP2719506) in compliance with UK Home Office Animals (Scientific Procedures) Act 1986 guidelines.

### Mice

Rosa26^lsl-tdTomato^ (B6.Cg-*Gt(ROSA)26Sor^tm^*^14^*^(CAG-tdTomato)Hze^*/J), Rosa26^lsl^-^iDTR^ (CBy.B6-*Gt(ROSA)26Sor^tm1(HBEGF)Awai^*/J) and transgenic Gzmb-Cre (B6;FVB-Tg(GZMB-cre)^1Jcb/J^) mice strains were purchased from Jackson Laboratories, USA. Gzmb^CreERT2^ was obtained from Dr. M. de la Roche, Cancer Research UK, Cambridge under MTA. *Gzmb^KI-Cre^* was generated by Ingenious Targeting Laboratory for Dr. Mahima Swamy. Mice were maintained in a standard barrier facility on a 12hr light/dark cycle at 21°C in individually ventilated cages with sizzler-nest material and fed an R&M3 diet (Special Diet Services, UK) and filtered water ad libitum. Cages were changed at least every 2 weeks. Age- and sex-matched mice between 8 and 20 weeks were used for all experiments. Mice of both genders were used unless otherwise indicated in the figure legends. No statistical methods were used to pre-determine sample size, but our sample sizes are similar to those reported in previous publications.

### Generation of *Gzmb^Cre-KI^* mice

*Gzmb^Cre-KI^* mice were made for MS by Ingenious Targeting Laboratories, USA. A 11.7 kb region used to construct the targeting vector was first sub cloned from a positively identified C57BL/6 fosmid clone (WI1-1012P9) using a homologous recombination-based technique. The region was designed such that 5’ homology arm extends 7.3 kb 5’ to exon 5 of the mouse *Gzmb* gene. The 2A-iCre-pA cassette was inserted immediately downstream of the stop codon (TAA) and replaced the 3’UTR sequence. An FRT-flanked hUbs-gb2 Neomycin cassette was inserted 343 bp downstream of the knockin cassette. The 3’ homology arm extended about 4.0 kb downstream from the Neo cassette. The targeting vector was linearized and transfected by electroporation of FLP C57Bl/6 (BF1) embryonic stem cells. After selection with G418 antibiotic, surviving clones were expanded for PCR analysis to identify recombinant ES clones. The Neo cassette in targeting vector was been removed during ES clone expansion. Targeted C57BL/6 FLP embryonic stem cells were microinjected into Balb/c blastocysts. Resulting chimeras with a high percentage black coat color were mated to C57BL/6 WT mice to generate Germline Neo deleted mice. Tail DNA from black mice were screened using multiple primers for Noe deletion, FLP absence, 5’ junction, and 3’ junction. One positive male mouse was backcrossed at least 5 generations to C57Bl/6J mice (Charles River UK), before crossing to other alleles. *Gzmb^Cre-KI^*developed in this study are available upon request with an MTA.

### Diphtheria Toxin administration for IEL depletion

To selectively deplete intraepithelial lymphocytes (IELs), IEL^WT^ and IEL^iDTR^ mice were administered a single intraperitoneal dose of diphtheria toxin (DT) at 50 μg/kg in a volume not exceeding 10 ml/kg. Tissues were harvested 24 h post-injection for downstream analysis or further experimental procedures.

### Isolation of IEL

IELs were isolated as previously described^42^. Briefly, the small intestine was removed and flushed with PBS. The intestine was cut longitudinally and then transversely into 0.5 cm pieces. Gut pieces were incubated in RPMI containing 10% FBS, Penicillin/Streptomycin, L-Glutamine and 1 mM DTT. After 40-minute incubation with shaking, pieces of gut were vortexed and filtered through a 100 μm cell strainer. Filtrate was spun and resuspended in 44% Percoll. This was layered on top of 75% Percoll (Sigma) and spun at 700g for 30 minutes without brake. Cells were removed from the interface of the Percoll layers and washed before further use. For single cell ATAC seq, IEL (Live, CD45+ CD3+ CD8a+) were flow sorted to purity of 99 percent and then pelleted and snap frozen for further processing.

### Lamina propria isolation

Small and large intestines were flushed with 20 ml of cold PBS and then longitudinally opened and stored on ice in 10 ml of PBS. Samples were vortexed 3 times for 10 s in PBS. Small intestines were then incubated for 30 min, with constant shaking, in “strip buffer” (PBS; 5 % FBS; 1 mM EDTA, 1 mM DTT), prewarmed at 37 °C. After the incubation, tissue was washed in PBS, then incubated for 45 min with constant shaking with “digest buffer” (RPMI; 10 % FBS; 1 mg/ml collagenase/dispase (Roche); 20 μg/ml DNAse 1 (Sigma). Supernatants containing LPL were then collected by pouring over a 70 μm filter into a new tube containing RPMI media supplemented with 10 % FBS.

### Isolation of lymphocytes from Peyer’s patches, liver, mesenteric lymph nodes, thymus and spleen

Organs were removed from mice and crushed through a 70 μm cell strainer into isolation media followed by RBC lysis. Except liver, cell suspensions of all other organs were resuspended in appropriate buffer for downstream applications. For liver, filtrate was spun and resuspended in 44% Percoll. This was layered on top of 75% Percoll (Sigma) and spun at 700g for 30 minutes without brake. Cells were removed from the interface of the Percoll layers and washed before further use.

### Tamoxifen preparation for Cre-ERT2 induction

Tamoxifen (Sigma-Aldrich, T5648) was dissolved in corn oil (Sigma-Aldrich, C8267) to a final concentration of 20 mg/ml. To prepare, 5 ml of corn oil was pre-warmed at 42 °C for 30 minutes. Tamoxifen powder, equilibrated to room temperature and protected from light, was weighed under a chemical hood and added to the warmed oil (100 mg per 5 ml). The solution was wrapped in foil and agitated at 37 °C on an orbital shaker (220 rpm) for 6–8 hours until fully dissolved. Vortexing was used intermittently to prevent clumping. The solution was stored at 4 °C, protected from light, and used within 7 days. Prior to administration, the solution was filtered using a 5 ml syringe fitted with an 18-gauge needle to prevent particulate injection. Tamoxifen was administered orally to minimize peritoneal inflammation associated with intraperitoneal delivery. In total, 4 consecutive intraperitoneal injections of 75 mg/kg were given on 4 consecutive days. 7 days post last injection was used as Day 1 for experiment.

### FTY720 treatment and IEL analysis

To assess the role of recirculating T cells in IEL repopulation, IEL^iDTR^ mice were administered a single intraperitoneal injection of diphtheria toxin (DT; 50 μg/kg, <10 ml/kg) on day 0 to deplete IEL. Beginning 24 hours post-DT (day 1), mice received daily intraperitoneal injections of FTY720 (1 mg/kg, <10 ml/kg; Cayman Chemical) or vehicle control (sterile water) for six consecutive days. Untreated littermates served as baseline controls. Mice were euthanized on day 7 for isolation and analysis of intestinal intraepithelial lymphocytes.

### EdU incorporation assay

To assess IEL proliferation, mice were injected intraperitoneally with 1.25 mg of 5-ethynyl-2’-deoxyuridine (EdU; Thermo Fisher Scientific) on day 3 post-diphtheria toxin (DT) administration. A fresh EdU stock solution was prepared by dissolving 50 mg of EdU in 6 ml of sterile PBS (final concentration: 8.33 mg/ml), with heating at 55 °C to aid dissolution. Each mouse received 150 µl of this solution (1.25 mg EdU) in a total injection volume of <10 ml/kg body weight. IEL were isolated as mentioned above, followed by EdU detection using the Click-iT™ Plus EdU Alexa Fluor™ 488 Imaging Kit (Thermo Fisher Scientific), according to manufacturer’s instructions.

### Flow Cytometry

The following murine monoclonal antibodies were used to detect cell surface markers: TCRβ [clone H57-597 (BioLegend)],TCRγδ[clone GL3 (BioLegendoreBioscience)], CD4 [clone RM4-5 (BioLegend)], CD8α [clone 53–6.7 (BioLegend)], CD8β [clone H35-17.2 (eBioscience)], CD103 [clone 2E7 (BioLegend)], CD45 [clone 30-F11 (BioLegend)], CD3 [clone 17A2 (BioLegend)]. Live dead stains used are – DAPI (Invitrogen, D1306), 7-AAD (Biolegend, 420403), near IR (Thermofisher, L10119), Ghost 450 (Cytek Biosciences, 13-0863). For intracellular staining, cells were fixed with 2 % PFA at 37 °C for 10 min before permeabilization with permabilization buffer (eBioscience). Cells were incubated with the following murine monoclonal antibodies: GzmB[clone GB12 (eBioscience)].

### Single cell ATAC sequencing

Frozen IEL were sent to ACTIV MOTIF in Belgium. Cells were harvested and frozen in culture media containing FBS and 10% DMSO. Cryopreserved cells were sent to Active Motif to perform the scATAC-seq assay. The cells were thawed in a 37°C water bath and prepared as described by 10X Genomics Demonstrated Protocol – Nuclei Isolation for Single Cell ATAC Sequencing Rev B. Briefly, cell pellets were resuspended in lysis buffer and incubated on ice for 5 minutes. Lysed cells were washed, strained, and nuclei were resuspended and counted using a Countess II FL Automated Cell Counter. Isolated nuclei were then used as input following the 10X Genomics Chromium Next GEM Single Cell ATAC Reagent Kits v1.1 manual. Targeting a 5,000 nuclei recovery, samples were added to the tagmentation reaction, loaded into the Chromium Controller for nuclei barcoding, and prepared for library construction following manufacturer’s protocol (10X Genomics PN-1000175). Resulting libraries were quantified using the KAPA Library Quantification Kit for Illumina platforms (KAPA Biosystems), and sequenced with PE34 sequencing on the NextSeq 500/550 sequencer (Illumina). Sequenced data were processed with the Cell Ranger ATAC software, with alignment to the mm10 genome. The Cell Ranger output files were used as input to Active Motif’s proprietary analysis program, which creates Excel tables containing detailed information on cluster-specific peak locations, gene annotations, and motif enrichment. The alignment files generated by Cell Ranger were also processed as pseudo-bulk ATAC-Seq samples. Duplicate reads were removed, only reads mapping as matched pairs and only uniquely mapped reads (mapping quality >= 1) were used for further analysis. Alignments were extended in silico at their 3’-ends to a length of 200 bp and assigned to 32-nt bins along the genome. The resulting histograms (genomic “signal maps”) were stored in bigWig files. Peaks were identified using the MACS 2.1.0 algorithm at a cutoff of p-value 1e-7, without control file, and with the –nomodel option. Peaks that were on the ENCODE blacklist of known false ChIP-Seq peaks were removed. Signal maps and peak locations were used as input data to Active Motif’s proprietary analysis program, which creates Excel tables containing detailed information on sample comparison, peak metrics, peak locations and gene annotations. MACS: Zhang et al. Model-based Analysis of ChIP-Seq (MACS). Genome Biol (2008) vol. 9 (9) pp. R137. <u>Other key software used:</u> Cell Ranger ATAC (v1.2.0) (demultiplexes BCL files, processes FASTQ files, and aggregates samples) Samtools (v0.1.19) (processing of BAM files), BEDtools (v2.25.0) (processing of BED files) and wigToBigWig (v4) (generation of bigWIG files)

### Immunofluorescence and H&E staining of small intestinal sections

Jejunal tissue was harvested from mice, flushed once with ice-cold HBSS and once with room-temperature 2% paraformaldehyde (PFA) in PBS (pH 7.4), and cut into ∼0.8–1.0 cm fragments. Tissues were fixed for 2–3 h at room temperature in 10 ml of fresh 2% PFA under gentle agitation. Fixed samples were washed three times in 50 mM ammonium chloride in PBS to quench residual aldehydes and incubated overnight in 30% sucrose (w/v) in PBS for cryoprotection. Tissues were then embedded in OCT compound (Agar Scientific), snap-frozen on dry ice, and stored at −18 °C or below. For Gzmb^cre/+^; tdTomato mice, cryosections were incubated with Phalloidin-FITC to label actin, and DAPI, before being imaged. Cryosections (15 μm) were rehydrated in PBS, permeabilized for 10– 15 min in 1% NP-40 in PBS, and blocked for 1 h at room temperature in PBS containing 2% BSA and 0.1% Triton X-100. Sections were incubated overnight at 4 °C with primary antibodies diluted in blocking buffer: goat anti-granzyme B (R&D Systems, AF1865; 1:50), mouse anti-E-cadherin (BD Transduction Laboratories, 610182; 1:150), and rabbit anti-CD3 (Dako/Agilent, A0452; 1:100). After five PBS washes, slides were incubated for 1 h at room temperature with the following secondary antibodies (1:500, Jackson ImmunoResearch): Alexa Fluor 488-conjugated donkey anti-goat, Alexa Fluor 647-conjugated donkey anti-mouse, and Alexa Fluor 568-conjugated goat anti-rabbit. Following four additional PBS washes, nuclei were counterstained with DAPI (1 μg/ml in PBS, 10 min), and sections were mounted using Vectashield Vibrance Antifade Mounting Medium (Vector Laboratories). Images were acquired using a Zeiss LSM 710 or LSM 880 confocal microscope with a 63×/1.4 NA oil-immersion objective, operated via Zen software. Where indicated, z-stacks spanning the full tissue depth were acquired and maximal intensity projections were generated using ImageJ.

For histological analysis, fixed jejunal samples were processed for paraffin embedding, sectioned at 5 μm, and stained with hematoxylin and eosin (H&E) using standard protocols. Images were acquired on a Zeiss Axioscan 7-2 slide scanner.

### RNA isolation, cDNA, and qPCR

Tissue samples were lysed in lysis buffer using 5 μm steel beads with the Qiagen TissueLyser LT, set to 50 oscillations for 2 minutes. Total RNA was extracted using the Purelink RNeasy Mini Kit (Thermo Fisher Scientific), following the manufacturer’s instructions. Subsequently, 1 μg of total RNA was reverse transcribed into cDNA using the PrimeScript™ 1st Strand cDNA Synthesis Kit (TaKaRa). For gene expression analysis, the resulting cDNA was combined with TAKARA TB Green® Premix Ex Taq™ II (Tli RNase H Plus) and gene-specific primer pairs listed in Table X. TBP served as the housekeeping gene. Quantitative real-time PCR (qRT-PCR) was performed on a BIO-RAD CFX96 C1000 Touch Real-Time PCR Detection System. Relative gene expression was determined using the ΔΔCt method and normalized to the housekeeping gene. All assays were run in technical triplicates.

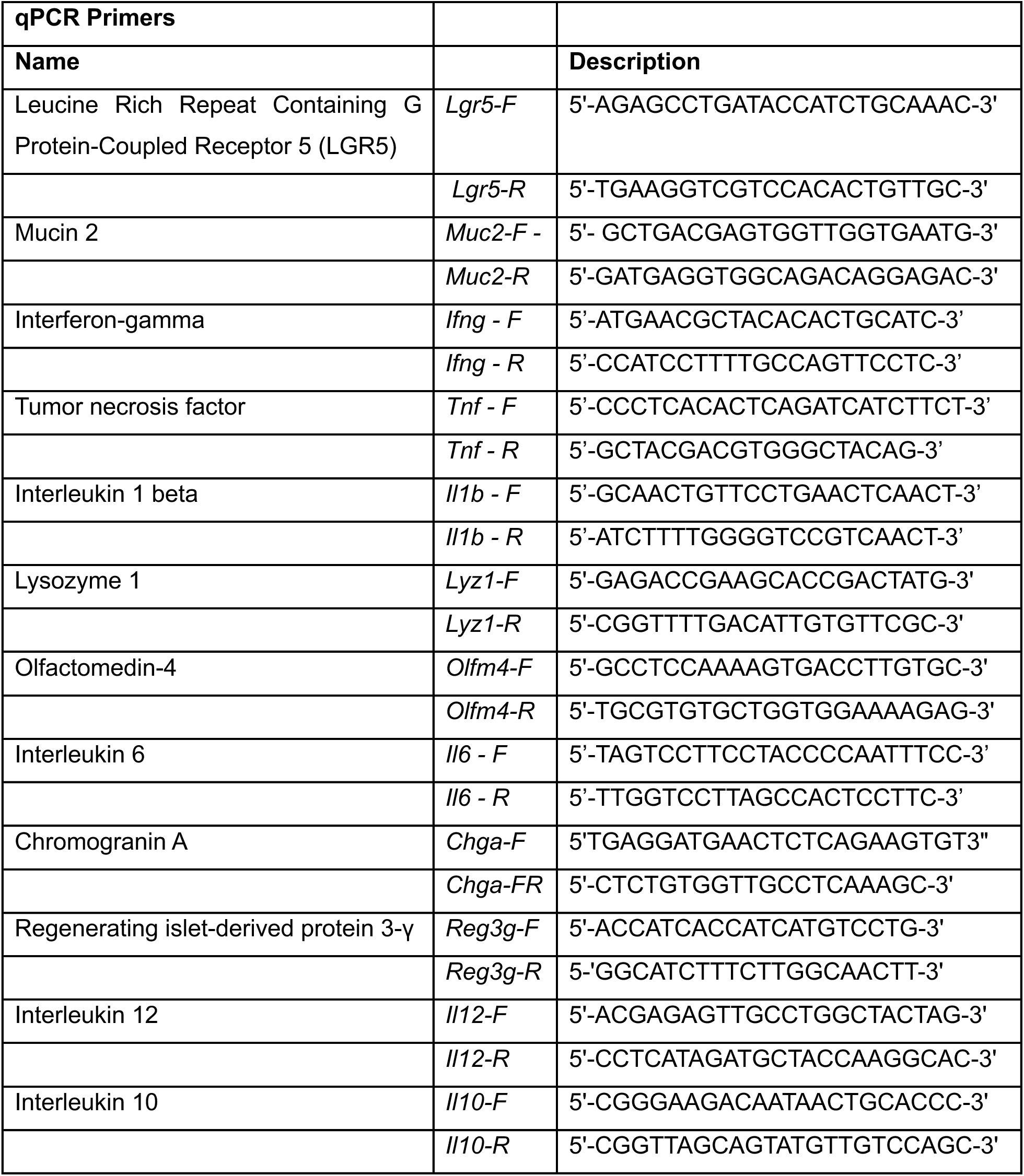
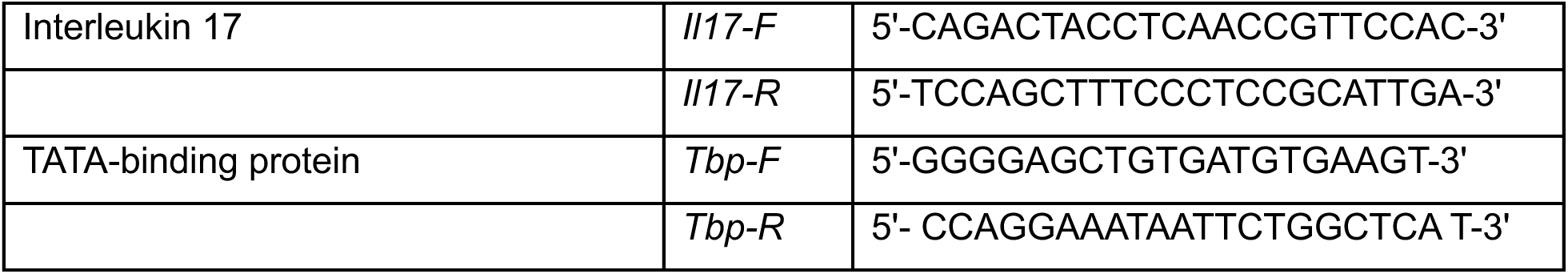

### Microbiota analysis

Small intestinal contents were collected from mice on day 2 post-diphtheria toxin (DT) administration. Total microbial DNA was extracted from these contents using the DNeasy PowerSoil Pro Kit (Qiagen, Cat. No. 47014), following the manufacturer’s protocol. Concentrations were normalized to 1 ng/μL using sterile distilled water and snap frozen and sent to Novogene for further processing. The V3–V4 hypervariable region of the 16S rRNA gene was PCR-amplified using universal primers 341F (5′-CCTAYGGGRBGCASCAG-3′) and 806R (5′-GGACTACNNGGGTATCTAAT-3′). PCR reactions (15 μL) were prepared using Phusion® High-Fidelity PCR Master Mix (NEB), primers (2 μM), and template DNA (∼10 ng). The cycling conditions were: 98 °C for 1 min, 30 cycles of 98 °C for 10 s, 50 °C for 30 s, 72 °C for 30 s, followed by a final extension at 72 °C for 5 min. Amplicons were confirmed on 2% agarose gels, pooled in equimolar amounts, and purified using the Qiagen Gel Extraction Kit. Libraries were constructed using the TruSeq® DNA PCR-Free Library Prep Kit and quality-checked using a Qubit® 2.0 Fluorometer and Agilent Bioanalyzer 2100. Sequencing was performed on an Illumina NovaSeq 6000 platform with 250 bp paired-end reads.

Bioinformatics Analysis: Raw paired-end reads were demultiplexed using unique sample barcodes, and primers and barcodes were trimmed. Reads were merged using FLASH v1.2.11, and quality-filtered using fastp v0.23.1. Chimeras were removed using the UCHIME algorithm with reference to the SILVA and UNITE databases.

ASV Denoising and Taxonomy Assignment: Denoising and ASV generation were performed using the DADA2 plugin in QIIME2 (v2022.2). Taxonomic annotation was done using the SILVA database for 16S rRNA gene sequences. Where species-level assignments were missing, taxonomy was corrected using the NCBI database. Phylogenetic relationships among ASVs were inferred through multiple sequence alignment in QIIME2.

Normalization and Community Composition: Sequence counts were normalized to the sample with the lowest read depth. Relative abundance bar plots (top 10 taxa) and heatmaps (top 35 taxa) were generated using Perl (SVG) and R (pheatmap), respectively. Ternary plots were constructed using the vcd() function in R. Venn and flower diagrams were used to show shared and unique taxa using R (VennDiagram) and Perl (SVG). A phylogenetic tree was generated for the top 100 genera using alignment and Perl visualization. Alpha and Beta Diversity Analysis: Diversity metrics including Observed_otus, Chao1, Shannon, Simpson, Dominance, good’s coverage, and Pielou_e were calculated using QIIME2. Species accumulation boxplots were created in R (vegan package) to assess sample richness and adequacy. Beta Diversity and Ordination: Beta diversity was assessed using weighted and unweighted UniFrac distances via QIIME2. Heatmaps of UniFrac distances were generated in Perl. UPGMA clustering was performed based on these distances. Ordination analyses including PCA, PCoA, and NMDS were conducted in R (ade4, ggplot2) to visualize community differences. Differential Abundance and Functional Prediction Statistical Testing: Community composition differences were evaluated using ANOSIM, ADONIS, MRPP, and SIMPER, implemented in R (vegan, ggplot2). Differential abundance was also assessed using MetaStat and LEfSe for biomarker discovery. Association Analysis: Species co-occurrence and correlations with environmental variables were analyzed via Spearman correlation, CCA, RDA, and dbRDA in R. 2D and 3D network visualizations were also generated. Software EnvironmentBioinformatics analyses were performed using Python v3.6.13, R v4.0.3 and Perl v5.26.2.

### Oral challenges with pathogens and chemicals

#### *Salmonella* typhimurium

*Salmonella enterica* serovar typhimurium SL1344 were grown overnight in LB broth supplemented with ampicillin (100 μg/ml) at 37 °C, on a shaker, then subcultured in LB+ampicillin medium for at least 3 h before *in vivo* infection. Bacteria were centrifuged 10 min at 3750 rpm, washed and resuspended in sterile PBS. The OD600 was measured to estimate bacterial density. Serial plating on LB agar supplemented with ampicillin (100 μg/ml) were performed to quantify the infection dose. IEL^WT^ and IEL^iDTR^ mice injected IP with 50 µg/kg DT 24 hours prior to infection. All mice fasted for 4 hours immediately prior to infection with wildtype SL1344 by oral gavage (1.5 x 10^8^ CFU) in 100 μl of PBS per mouse (Refer Our mucosal immunology paper). For output cfu determination, mice were euthanized by a rising concentration of CO_2_ and death was confirmed by cervical dislocation. Fecal content, mLNs and spleen were collected, weighed, and transferred into 2 ml Precellys (Bertin) tubes containing ceramic beads and PBS+0.05 % Triton X-100 (Sigma). Tissues were homogenized using the Precellys 24 homogenizer (Bertin) for 2x10s at 5000 Hz. Livers were weighed and crushed through a 70 μm strainer in PBS+0.05 % Triton X-100. Supernatants were serial diluted in LB+ampicillin (100 μg/ml) medium and plated on LB+ampicillin plates.

#### *Listeria Monocytogenes* Infection

A murinized, lux-expressing strain of *Listeria monocytogenes* (Lmo-InA^m^-Lux; obtained from Andreas was streaked on BHI agar and grown at 37 °C. A single colony was used to inoculate 5 ml BHI broth, and cultures were incubated overnight at 37 °C with shaking. The following morning, bacterial density was measured by OD600 (typically ∼1.31), and cultures were diluted to an OD600 of 0.05 in fresh BHI for subculturing. At mid-log phase (∼4 h), the OD600 was rechecked and used to estimate bacterial density. Cultures were pelleted (3750 rpm, 10 min), washed three times in sterile PBS, and resuspended in PBS to achieve a final concentration of ∼2.5 × 10¹⁰ CFU/ml.

To confirm infectious dose, serial dilutions of the inoculum were plated on BHI agar, and colonies were counted the next day. Mice were fasted for 4 h prior to infection. Thirty minutes before infection, mice received 50 µl of 7.5% sodium bicarbonate in HBSS by oral gavage to neutralize stomach pH. Mice were then infected with 200 µl of the prepared Listeria suspension (∼2.7 × 10⁹ CFU) by oral gavage.

For output CFU analysis, mice were euthanized by a rising concentration of CO₂ followed by cervical dislocation. Small intestine (ileum), mesenteric lymph nodes (mLNs), spleen, and liver were collected, weighed, and homogenized in 2 ml PBS with 0.05% Triton X-100 using a Precellys 24 tissue homogenizer (Bertin) in ceramic bead tubes. Homogenates were serially diluted in LB broth and plated on BHI agar for bacterial enumeration.

#### *Cryptosporidium Parvum* infection

Mice injected IP with 50 µg/kg DT 24 hours prior to infection. All mice fasted for 4 hours immediately prior to infection with C. parvum by oral gavage (50,000 oocysts). Strain of C. parvum genetically modified to express NanoLuciferase as previously described (Hanna et al 2023, PMID: 37600947). Fecal samples collected daily starting day 3 post infection until day 9 post infection. Fecal samples were collected and analysed both at the individual animal and cage level for NanoLuciferase activity as previously described (Hanna et al 2023, PMID: 37600947).

#### *H. polygyrus* infection

Mice injected IP with 50 µg/kg DT 24 hours prior to infection. Mice were infected with 200 *H. polygyrus bakeri* L3 larvae via oral gavage and fecal pellets were collected through the infection at days 14, 21 and 28 for egg counts. Egg counts were carried out following weighing of the pellet(s) and resuspending in 1-2 mL of water overnight, vortexed to create a slurry and then adding an equal volume of a saturated salt solution. Egg counts were calculated via counting both sides of a McMaster slide and averaged, then a final count calculated considering dry fecal pellet weight. Mice were sacrificed 28 days post infection for measurement of adult worm burden, whereby adult worms were manually counted under a dissection microscope, and fecundity was calculated dividing the number of eggs by the number of worms counted at 28 days post infection.

#### Dextran Sulfate Sodium (DSS)-Induced Colitis Model

IEL^WT^ and IEL^iDTR^ mice injected IP with 50 µg/kg diptheria toxin 24 hours prior to infection. To induce colitis, mice were administered 3% (w/v) dextran sulfate sodium (DSS; MW 36,000–50,000 Da, MP Biomedicals) in drinking water *ad libitum* for 6 consecutive days. Mice were weighed and monitored daily. Clinical disease severity was assessed using a composite Disease Activity Index (DAI), incorporating weight loss, coat condition, and activity level:

| Scoring Sheet |  |  |  |
| --- | --- | --- | --- |
| Score | weight loss % | Coat condition | Level of Activity |
| 0 | 0 | good | active |
| 1 | 1-5% | slightly ruffled | active but less so than control |
| 2 | 5-10% | ruffled | slower |
| 3 | 10-15% | clear piloerection | slow, tentative |
| 4 | 15-20% | piloerection & hunched | inactive |
| 5 | >20% | piloerection & hunched & wasting | inactive when provoked |

In addition, as presence of blood in stools is an indicator of disease severity, we will also independently monitor blood in the stools using the following grading system:

0: Normal Stools

1: Soft or very soft stools

2: Very soft stools with visual traces of blood 3: Watery stools with visible rectal bleeding

A score of 3 for the faecal scoring system will result in the animal(s) being culled.

### Data analysis

Flow cytometry data was analyzed using FlowJo Software (v10.9.0). Data analysis and statistical tests were carried out using GraphPad Prism (v10.0.0).

---

**Supplementary Figure 1.**
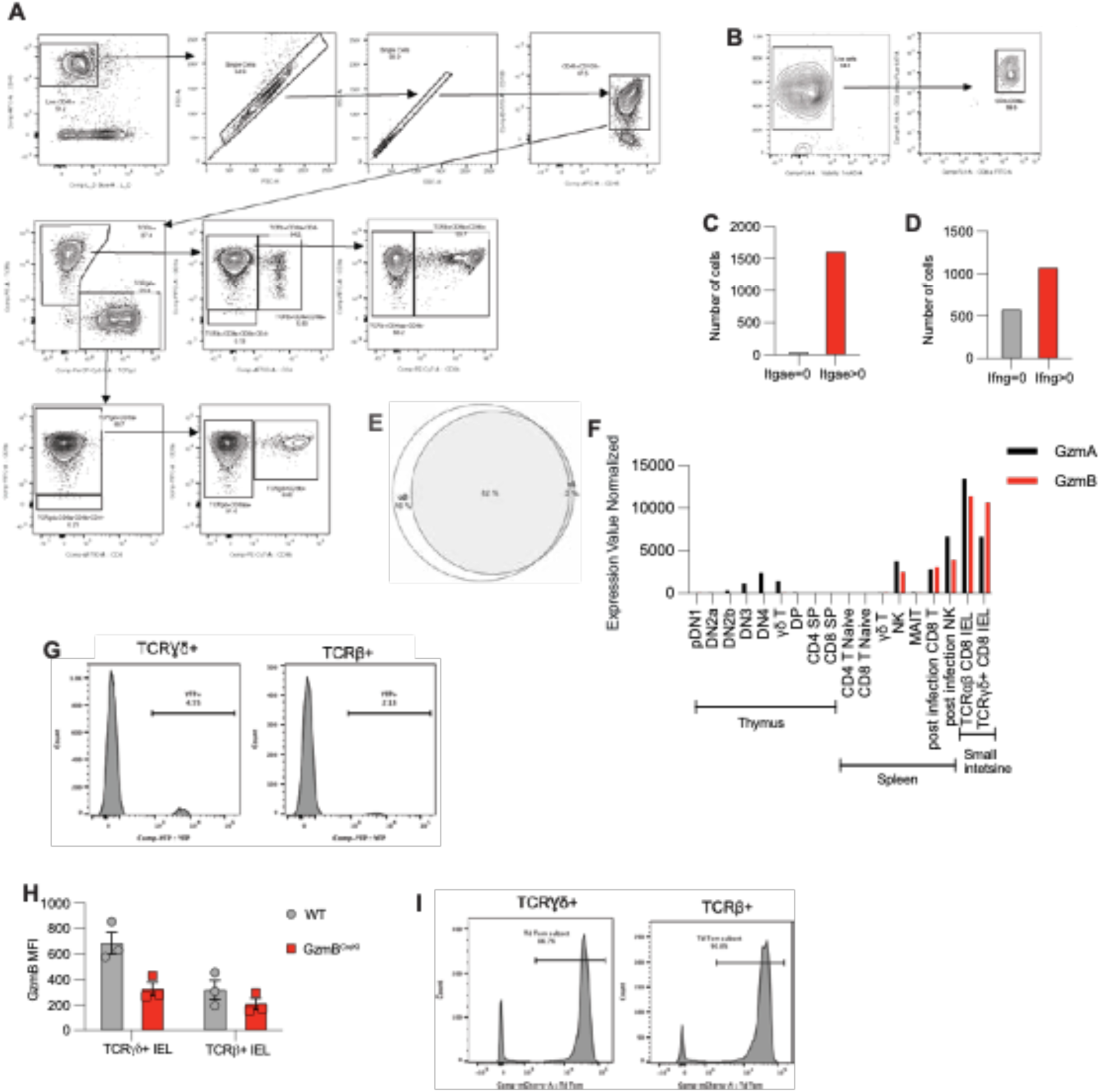
**A**) Flow cytometry gating strategy used to define IEL subsets from small intestinal epithelial preparations. **B)** Sorting strategy for scATAC-seq analysis. Live CD45⁺CD3⁺CD8α⁺ IEL were isolated for chromatin profiling. **C–D)** Bar graphs quantifying the proportion of IELs with accessible chromatin at (C) *Itgae* (encoding CD103) and (D) *Ifng*, canonical markers of IEL identity and effector function. **E)** Area-proportional Venn diagram illustrating the overlap in accessible chromatin regions between TCRγδ⁺ and TCRαβ⁺ IEL based on previously published bulk ATAC-seq data (Semenkovich et al., 2016), indicating extensive epigenetic convergence. **F)** Normalized RNA expression of *Gzma* and *Gzmb* across T cell developmental stages, mature T cell subsets, and NK cells, based on publicly available ImmGen datasets (https://www.immgen.org/). **G)** Representative histogram showing inefficient IEL labeling in previously reported *GzmbCre* transgenic mice crossed to *Rosa26*^lsl-EYFP^. **H)** Representative histogram of tdTomato⁺ IEL in *Gzmb*^Cre/+^× *Rosa26*^lsl-tdTomato^ mice, demonstrating robust and specific labeling across IEL subsets. **I)** Flow cytometry quantification of intracellular GzmB protein expression across major IEL subsets from *Gzmb*^Cre/+^mice.

**Supplementary Figure 2.**
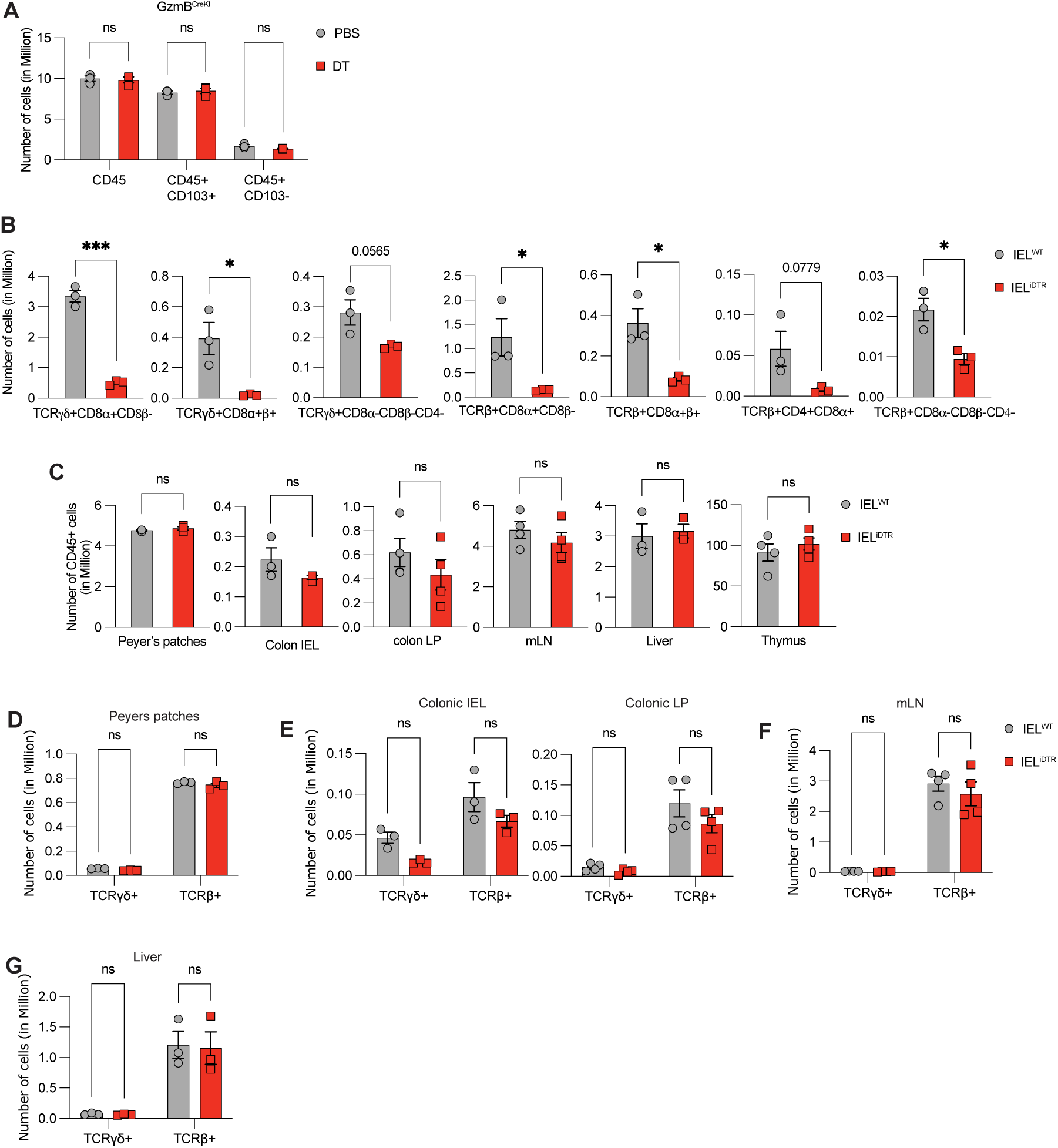
**A**) Flow cytometric quantification of CD45⁺, CD45⁺CD103⁺ (IEL), and CD45⁺CD103⁻ (non-IEL) subsets in the small intestine of *Gzmb*^Cre/+^ mice 24 hours after diphtheria toxin (DT) or PBS administration. Data represent mean ± SEM from 3 mice per group. **B)** Quantification of individual IEL subsets in the small intestine of IEL^WT^ and IEL^iDTR^ mice at day 1 post-DT, confirming broad depletion across γδ and αβ IEL lineages. Data represent mean ± SEM from 3 mice per group. **C)** Flow cytometric analysis of total CD45⁺ immune cells in multiple peripheral and mucosal compartments, including Peyer’s patches, colon IEL, colon lamina propria (LP), mesenteric lymph nodes (mLN), liver, and thymus of IEL^WT^ and IEL^iDTR^ mice, 24 hours after DT treatment. Data represent mean ± SEM from 3 mice per group. **D–G)** Quantification of TCRγδ⁺ and TCRβ⁺ lymphocyte subsets across mucosal and systemic tissues—Peyer’s patches, colon IEL, colon LP, mLN, and liver of IEL^WT^ and IEL^iDTR^ mice —24 hours post-DT administration. Data represent mean ± SEM from 3 mice per group. Statistical comparisons: ANOVA with Sidak’s post hoc correction (A, D, E, F, G) and unpaired t-test (B, C). **\*** p < 0.05, ** p < 0.01, *** p < 0.001, ns = non-significant.

**Supplementary Figure 3.**
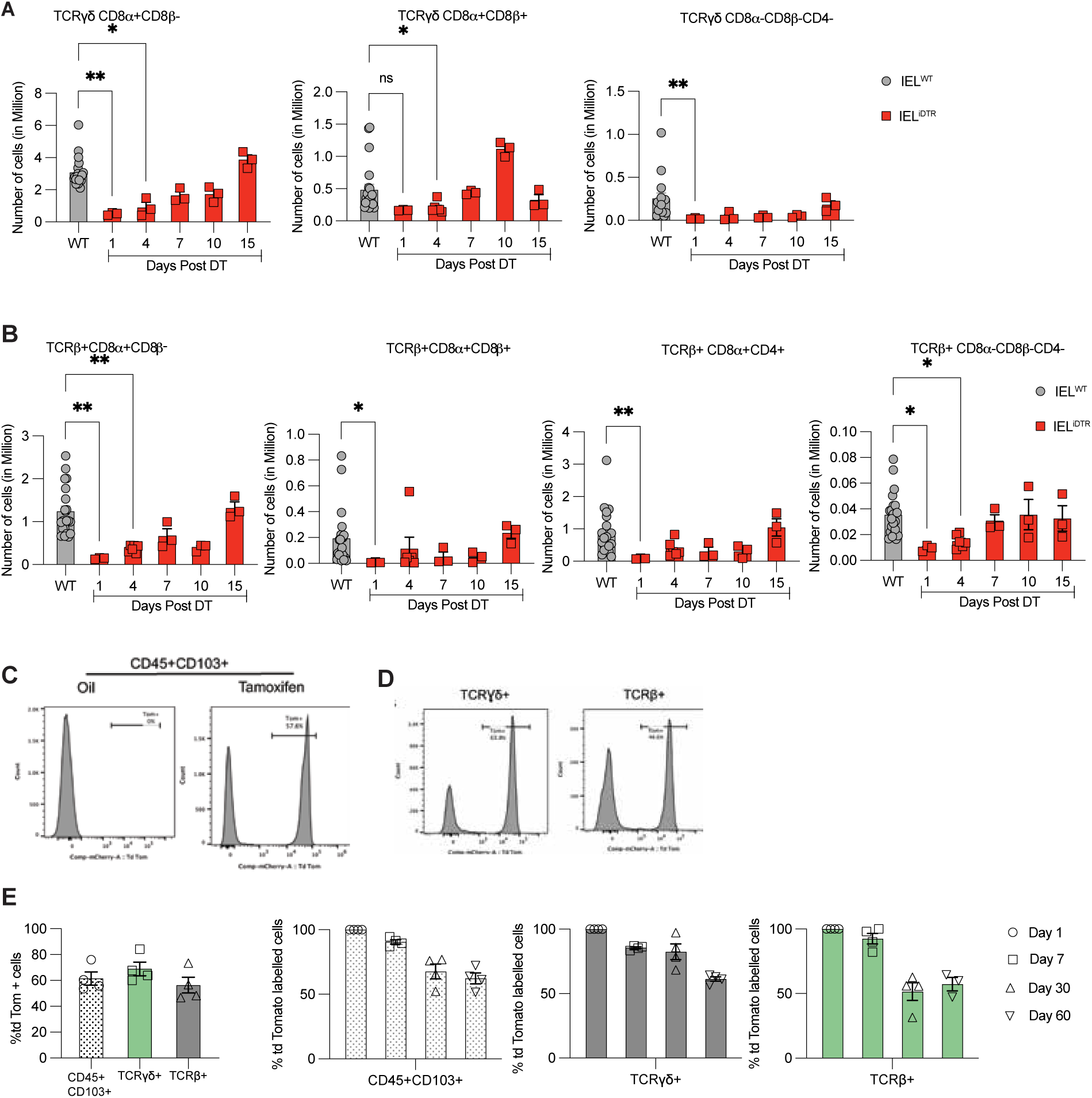
**A–B**) Time-course quantification of (A) TCRγδ⁺ and (B) TCRβ⁺ IEL subsets following a single DT injection in IEL^WT^ and IEL^iDTR^ mice, demonstrating progressive repopulation over 15 days. Data represent mean ± SEM from 3-5 mice per group. **C–D)** Representative histograms of tdTomato expression in (D) CD45⁺CD103⁺ and (E) TCRγδ⁺ and TCRβ⁺ IEL from *Gzmb*^CreERT2^ × *Rosa26*^lsl-tdTomato^ mice, assessed 7 days after final tamoxifen or oil-only control administration. **E)** Quantification of tdTomato⁺ IEL across indicated time points, demonstrating the longevity of labelled IEL subsets. Data represent mean ± SEM from 4 mice per group. Statistical comparisons: ANOVA with Sidak’s post hoc correction (A, B). **\*** p < 0.05, ** p < 0.01.

